# Castration delays epigenetic aging and feminises DNA methylation at androgen-regulated loci

**DOI:** 10.1101/2020.11.16.385369

**Authors:** VJ Sugrue, JA Zoller, P Narayan, AT Lu, OJ Ortega-Recalde, MJ Grant, CS Bawden, SR Rudiger, A Haghani, DM Bond, M Garratt, KE Sears, N Wang, XW Yang, RG Snell, TA Hore, S Horvath

**Affiliations:** Department of Anatomy, University of Otago, Dunedin, 9016, New Zealand; Department of Biostatistics, Fielding School of Public Health, University of California Los Angeles, Los Angeles, California, USA; Applied Translational Genetics Group, School of Biological Sciences, Centre for Brain Research, The University of Auckland, Auckland, 1010, New Zealand; Department of Human Genetics, David Geffen School of Medicine, University of California Los Angeles, Los Angeles, CA 90095, USA; Livestock and Farming Systems, South Australian Research and Development Institute, Roseworthy, South Australia 5371, Australia; Department of Ecology and Evolutionary Biology, UCLA, Los Angeles, CA, USA; Center for Neurobehavioral Genetics, Semel Institute for Neuroscience and Human Behavior, University of California, Los Angeles (UCLA), Los Angeles, CA 90095, USA; Department of Psychiatry and Biobehavioral Sciences, David Geffen School of Medicine at UCLA, Los Angeles, CA 90095, USA

**Keywords:** Castration, epigenetic clock, epigenetics, DNA methylation, aging/ageing, biomarker of aging, androgens

## Abstract

In mammals, females generally live longer than males. Nevertheless, the mechanisms underpinning sex-dependent longevity are currently unclear. Epigenetic clocks are powerful biological biomarkers capable of precisely estimating chronological age using only DNA methylation data. These clocks have been used to identify novel factors influencing the aging rate, but few studies have examined the performance of epigenetic clocks in divergent mammalian species. In this study, we developed the first epigenetic clock for domesticated sheep (*Ovis aries*), and using 185 CpG sites can predict chronological age with a median absolute error of 5.1 months from ear punch and blood samples. We have discovered that castrated male sheep have a decelerated aging rate compared to intact males, mediated at least in part by the removal of androgens. Furthermore, we identified several androgen-sensitive CpG dinucleotides that become progressively hypomethylated with age in intact males, but remain stable in castrated males and females. Many of these androgen sensitive demethylating sites are regulatory in nature and located in genes with known androgen-dependent regulation, such as *MKLN1, LMO4* and *FN1*. Comparable sex-specific methylation differences in *MKLN1* also exist in mouse muscle (p=0.003) but not blood, indicating that androgen dependent demethylation exists in multiple mammalian groups, in a tissue-specific manner. In characterising these sites, we identify biologically plausible mechanisms explaining how androgens drive male-accelerated aging.

## INTRODUCTION

Age has a profound effect on DNA methylation in many tissues and cell types (Horvath, 2013; Issa, 2014; Rakyan et al., 2010; Teschendorff et al., 2010). When highly correlated age-dependent sites are used as a model through the use of a tool known as the epigenetic clock, exceptionally precise estimates of chronological age (termed “DNAm age” or “epigenetic age”) can be achieved using only purified DNA as an input (Hannum et al., 2013; Horvath, 2013; Horvath and Raj, 2018; Levine et al., 2018). For example, despite being one the earliest epigenetic clocks constructed, Horvath’s 353 CpG site clock is capable of estimating chronological age with a median absolute error (MAE) of 3.6 years and an age correlation of 0.96, irrespective of tissue or cell type (Horvath, 2013). Estimates generated by this and related epigenetic clocks are not only predictive of chronological age but also biological age, allowing identification of pathologies as well as novel genetic and environmental factors that accelerate or slow biological aging. For example, irrespective of ethnic background, females and exceptionally long-lived individuals are found to have reduced epigenetic aging compared to males and other controls (Horvath et al., 2015, 2016).

Lifespan in mammals (including humans) is highly dependent upon an individual’s sex, whereby females generally possess a longevity advantage over males (Lemaître et al., 2020). Despite being a fundamental risk factor affecting age-related pathologies, the mechanistic basis of how sex influences aging is relatively unexplored. Perhaps not surprisingly, sex hormones are predicted to play a central role, with both androgens and estrogens thought to influence aspects of the aging process (Horstman et al., 2012). Castration has been shown to extend the lifespan of laboratory rodents (Asdell et al., 1967), as well as domesticated cats (Hamilton, 1974) and dogs (Hoffman et al., 2013). Castration has also been associated with longer lifespans in historical survival reports of 14^th^-20^th^ century Korean eunuchs (Min et al., 2012) and men housed in US mental institutions in the 20^th^ century (Hamilton and Mestler, 1969), although not in castrato opera singers, somewhat common in the 15^th^-19^th^ centuries (Nieschlag et al., 1993). Conversely, estrogen production appears to have some protective effect on aging in females, with ovariectomised mice having a shortened lifespan (Benedusi et al., 2015) and replacement of ovaries from young animals into old female mice extending lifespan (Cargill et al., 2003). Indeed, ovariectomy has been shown to accelerate the epigenetic clock (Stubbs et al., 2017), supporting predictions that estrogen production slows the intrinsic rate of aging relative to males. In humans, natural and surgical induction of menopause also hastens the pace of the epigenetic clock, while menopausal hormone therapy decreases epigenetic aging as observed in buccal cell samples (Levine et al., 2016). Female breast tissue sourced DNA used for epigenetic clock calculation has been found to be substantially older as determined by this method than any other sources of DNA (Horvath, 2013; Sehl et al., 2017), further indicating a link between sex hormones and epigenetic aging. However, the effects of castration and/or testosterone production on the epigenetic predictors of aging in males remained unknown in either humans or animal models prior to the current study.

Domesticated sheep (*Ovis aries*) represent a valuable, albeit underappreciated, large animal model for human disease and share with humans more similar anatomy, physiology, body size, genetics, and reproductive lifestyle as compared with commonly studied rodents (Pinnapureddy et al., 2015). With respect to aging, sheep exhibit a remarkable female-specific lifespan advantage (Lemaître et al., 2020), and Soay sheep of the Outer Hebrides represent a cornerstone research paradigm for longevity in wild mammal populations (Jewell, 1997). Moreover, sheep are extensively farmed (and males castrated) in many countries, allowing incidental study of the effect of sex and sex hormones in aging to occur without increasing experimental animal use (Russell and Burch, 1959).

Here we present the first sheep epigenetic clock and quantify its median error to 5.1 months, ∼3.5-4.2 % of their expected lifespan. Significantly, we found not only that castration affects the epigenome, but that the methylomes of castrated male sheep show reduced epigenetic aging compared to intact male and female counterparts, a result consistent with the increased longevity of castrated Soay sheep (Jewell, 1997). Many genomic regions and associated genes with differential age association between castrated and intact males were identified, some of which are known to be regulated or bound by androgen receptor (AR) in humans. Taken together, these findings provide a credible mechanistic link between levels of sex hormones and sex-dependent aging.

## METHODS

### DNA extraction and quantitation

Sheep DNA samples for this study were derived from two distinct tissues from two strains: ear tissue from New Zealand Merino, and blood from South Australian Merino.

### Sheep ear DNA source

Ear tissue was obtained from females and both intact and castrated male Merino sheep during routine on-farm ear tagging procedures in Central Otago, New Zealand. As a small piece of tissue is removed during the ear tagging process that is usually discarded by the farmer, we were able to source tissue and record the year of birth without altering animal experience, in accordance with the New Zealand Animal Welfare Act (1999) and the National Animal Ethics Advisory Committee (NAEAC) Occasional Paper No 2 (Carsons, 1998). The exact date of birth for each sheep is unknown, however, this was estimated to be the 18th of October each year, according to the date at which rams were put out with ewes (May 10th of each year), a predicted mean latency until mating of 12 days, and the mean gestation period from a range of sheep breeds (149 days) (Fogarty et al., 2005). Castration was performed by the farmer using the rubber ring method within approximately 5-50 days from birth as per conventional farming practice (National Animal Welfare Advisory Comittee, 2018). Mass of yearlings was recorded by the farmer for both castrated and intact male sheep at 6.5 months of age, as a part of routine growth assessment. In total, ear tissue from 138 female sheep aged 1 month to 9.1 years and 126 male sheep (63 intact, 63 castrates) aged 6 months to 5.8 years was collected and subjected to DNA extraction (Figure 1A).

**Figure 1.**
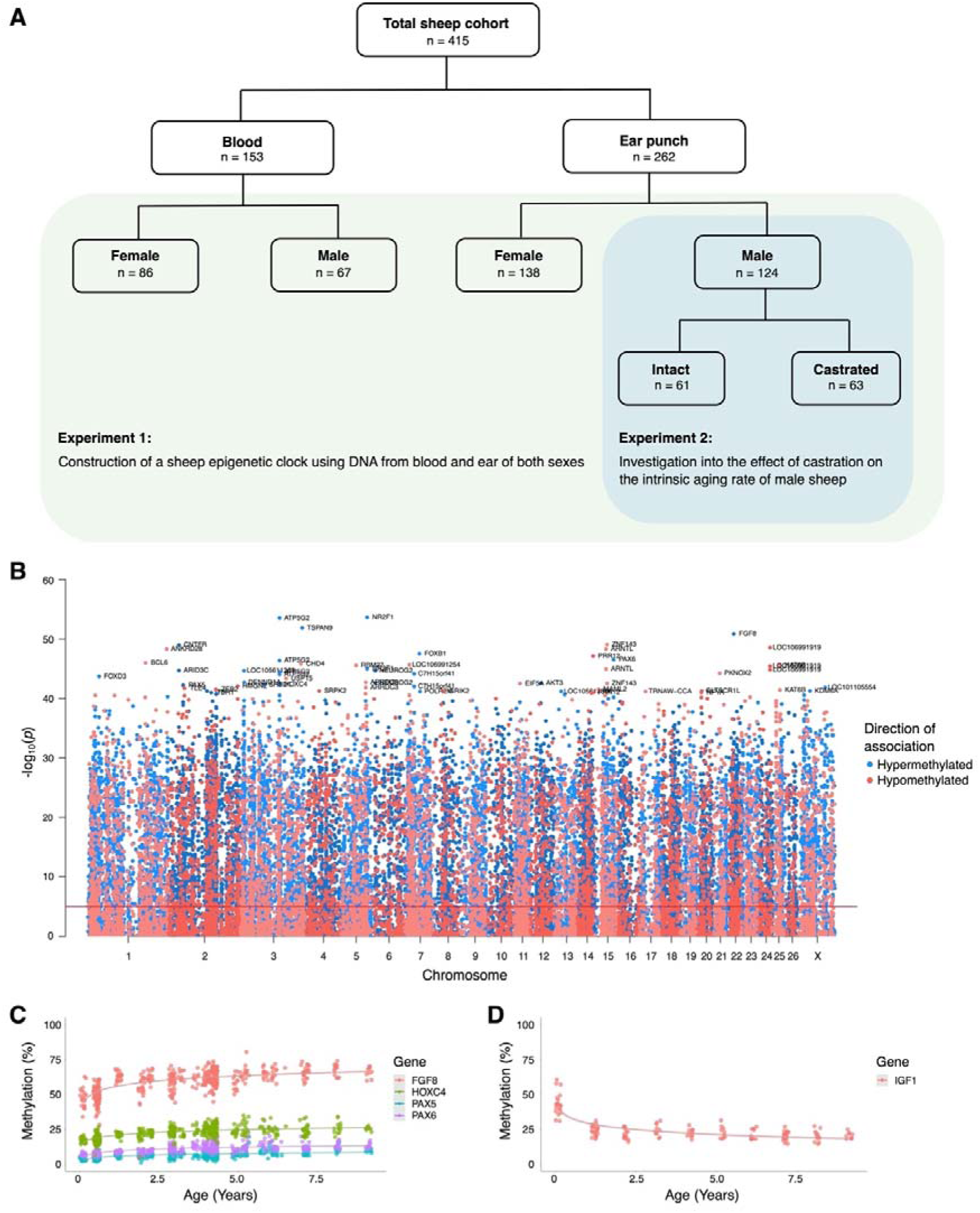
Creation of the epigenetic clock in sheep. **A)** Description of sheep cohort for this study. **B)** Manhattan plot of all CpGs and their correlation with chronological age. **C)** Methylation levels of highly age correlated probes within biologically relevant genes: *FGF8* cg10708287 (r=0.64, p=1.38E^-51^), *PAX6* cg00953859 (r=0.62, p=2.71E^-47^), *PAX5* cg16071226 (r=0.59, p=5.75E^-43^), *HOXC4* cg12097121 (r=0.59, p=4.47E^-43^). **D**) Methylation levels of *IGF1* cg18266944 in ear of females only (r=0.60, p=7.43^-15^). The *p* values of the correlation were calculated using the *standardScreeningNumericTrait* function in WGNCA (Student t-test).

DNA was extracted from ear punch tissue using a Bio-On-Magnetic-Beads (BOMB) protocol (Oberacker et al., 2019) which isolates DNA molecules using solid-phase reversible immobilisation (SPRI) beads. Approximately 3 mm punches of ear tissue were lysed in 200 μL TNES buffer (100 mM Tris, 25 mM NaCl, 10 mM EDTA, 10% w/v SDS), supplemented with 5 μL 20 mg/mL Proteinase K and 2 μL RNAse A and incubated overnight at 55 °C as per BOMB protocols. The remainder of the protocol was appropriately scaled to maximise DNA output while maintaining the necessary 2:3:4 ratio of beads:lysate:isopropanol. As such, 40 μL cell lysate, 80 μL 1.5X GITC (guanidinium thiocyanate), 40 μL TE-diluted Sera-Mag Magnetic SpeedBeads (GE Healthcare, GEHE45152105050250) and 80 μL isopropanol were combined. After allowing DNA to bind the SPRI beads, tubes were placed on a neodymium magnetic rack for ∼5 minutes until the solution clarified and supernatant was removed. Beads were washed 1x with isopropanol and 2x with 70% ethanol, and then left to air dry on the magnetic rack. 25 μL of MilliQ H_2_O was added to resuspend beads, and tubes were removed from the rack to allow DNA elution. Tubes were once again set onto the magnets, and the clarified solution (containing DNA) was collected.

DNA was quantified using the Quant-iT PicoGreen dsDNA assay kit (ThermoFisher Scientific, cat # P11496). 1 μL DNA sample was added to 14 μL TE diluted PicoGreen in MicroAmp optical 96-well plates (ThermoFisher Scientific, cat #N8010560) as per manufacturer directions, sealed, and placed into a QuantStudio qPCR machine for analysis. Samples with DNA content greater than the target quantity of 25 ng/μL were diluted with MilliQ.

### Sheep blood DNA source

DNA methylation was analysed in DNA extracted from the blood of 153 South Australian Merino sheep samples (80 transgenic Huntington’s disease model sheep (OVT73 line) (Jacobsen et al., 2010) and 73 age-matched controls) aged from 2.9 to 7.0 years (Figure 1A). All protocols involving OVT73 sheep were approved by the Primary Industries and Regions South Australia (PIRSA, Approval number 19/02) Animal Ethics Committee with oversight from the University of Auckland Animal Ethics Committee. The epigenetic age of the transgenic sheep carrying the *HTT* gene was not significantly different from controls (p=0.30, Mann-Whitney U test), therefore the data derived from these animals was subsequently treated as one dataset (Figure S1).

300 μL thawed blood samples were treated with 2 rounds of red cell lysis buffer (300 mM Sucrose, 5 mM MgCl_2_, 10 mM Tris pH8, 1% Triton X-100) for 10 minutes on ice, 10 minute centrifugation at 1,800 RCF, and supernatant removed between each buffer treatment. The resulting cell pellet was incubated in cell digestion buffer (2.4 mM EDTA, 75 mM NaCl, 0.5 % SDS) and Proteinase K (500 μg/ml) at 50 °C for two hours. Phenol:Chloroform:Isoamyl alcohol (PCI, 25:24:1; pH8) was added at equal volumes, mixed by inversion, and placed in the centrifuge for 5 minutes at 14,000 RPM at room temperature (repeated if necessary). The supernatant was collected and combined with 100% ethanol at 2x volume, allowing precipitation of DNA. Ethanol was removed and evaporated, and 50 μL TE buffer (pH8) was added to resuspend genomic DNA. DNA sample concentration was initially quantified using a nanodrop, followed by Qubit.

### Data processing and clock construction

A custom Illumina methylation array (“HorvathMammalMethyl40”) was used to measure DNA methylation. These arrays include 36k CpG sites conserved across mammalian species, though not all probes are expected to map to every species. Using QuasR (Gaidatzis et al., 2015), 33,136 probes were assigned genomic coordinates for sheep genome assembly OviAri4.

Raw .idat files were processed using the *minfi* package for RStudio (v3.6.0) with *noob* background correction (Aryee et al., 2014; Triche Jr et al., 2013). This generates normalised beta values which represent the methylation levels at probes on a scale between 0 (completely unmethylated) and 1 (fully methylated).

185 CpG sites were selected for the sheep epigenetic clock by elastic net regression using the RStudio package *glmnet* (Friedman et al., 2009). The elastic net is a penalized regression model which combines aspects of both ridge and lasso regression to select a subset of CpGs that are most predictive of chronological age. 88 and 97 of these sites correlated positively and negative with age, respectively. Epigenetic age acceleration was defined as residual resulting from regressing DNAm age on chronological age. By definition, epigenetic age acceleration has zero correlation with chronological age. Statistical significance of the difference in epigenetic age acceleration between each male group (castrated versus intact) was determined using a non-parametric two-tailed Mann-Whitney U test applied to sexually mature sheep only (> 18 months of age).

The human & sheep dual-species clock was created by combining our sheep blood and ear sourced data with human methylation data previously measured using the same methylation array (“HorvathMammalMethyl40”) (Horvath et al., 2020). This data comprises 1,848 human samples aged 0 to 93 years and includes 16 different tissues. The clock was constructed identically to the sheep only clock, with an additional age parameter *relative age* defined as the ratio of chronological age by maximum age for the respective species. The maximum age for sheep and humans was set at 22.8 years and 122.5 years, respectively, as defined in the anAge database (De Magalhães et al., 2009).

### Identification of age-associated and androgen-sensitive DMPs

Age-associated differentially methylated probes (DMPs) were identified using the weighted gene co-expression network analysis (WGCNA) function *standardScreeningNumericTrait* (Langfelder and Horvath, 2008) which calculates the correlation between probe methylation and age. Where mapped, gene names for the top 500 positively correlated probes were input into DAVID (Dennis et al., 2003) for functional classification analysis with the *Ovis aries* genes present on the methylation array as background. Androgen-sensitive DMPs (asDMPs) were identified using a t-test of the difference between the slopes of linear regression lines applied to methylation levels across age in each sex. A difference in slope indicates that there is an interaction between age and sex for methylation status of a particular probe.

### Transcription factor binding analysis

Transcription factor (TF) binding at asDMPs was evaluated by entering the equivalent human probe position into the *interval search* function of the Cistrome Data Browser Toolkit, an extensive online collection of chromatin immunoprecipitation sequencing (ChIP-seq) data (Mei et al., 2016; Zheng et al., 2019). TFs binding CpGs of interest in human were analysed using a custom R script, with ChIP-seq tracks being viewed by Cistrome link in the UCSC genome browser alongside additional annotation tracks of interest for export and figure creation in Inkscape. To model the background levels of TF binding, 1000 replicates of 50 random probes sites were run in a similar manner using the Cistrome Human Factor dataset, BEDtools (Quinlan and Hall, 2010) and a custom Python script.

## RESULTS

### DNA methylation in blood and ear throughout sheep aging

To create an epigenetic clock for sheep, we purified DNA from a total of 432 sheep of the Merino breed (Figure 1A). The majority of DNA samples (262) were from ear punches sourced from commercial farms in New Zealand, with the remaining (168) blood samples from a South Australian Merino flock. DNA methylation was quantified using a custom 38K probe array consisting of CpG sites conserved amongst a wide range of mammalian species; with 33,136 of these predicted to be complementary to sheep sequences. Two ear samples from intact males were excluded by quality control measures.

To initially characterise methylation data, we performed hierarchical clustering that revealed two major clusters based on tissue source (Figure S2A). There was some sub-clustering based on sex and age; however, there was no separation based on known underlying pedigree variation. Global average CpG methylation levels in ear tissue exhibited a small progressive increase with age, though the same trend was not seen in blood (Figure S2B).

Pearson correlation coefficients (r) describing the linear relationship between CpG methylation and chronological age ranged from -0.63 to 0.68 for all ear and blood samples (Table S1). One of the most positively correlated mapped probes was located within the promoter of fibroblast growth factor 8 (*FGF8*) (Figure 1C), a well-described developmental growth factor (r=0.64, p=1.38E^-51^). Probes located within several other well-known transcription factors were also among those most highly correlated with age (*PAX6*, r=0.62, p=2.71E^-47^; *PAX5*, r=0.59, p=5.75E^-43^; *HOXC4*, r=0.59, p=4.47E^-43^). Indeed, when we performed ontogeny analysis, we found the top 500 CpGs positively correlated with age were enriched for transcription-related and DNA binding processes (Table S2), consistent with wide-spread transcriptional shifts during different life stages. Interestingly, we also found a CpG (cg18266944) in the second intron of insulin-like growth factor 1 (*IGF1*) that becomes rapidly hypomethylated following birth before levelling off post-adolescence (r=-0.60, p=7.43E^-15^). We considered this a particularly encouraging age-associated epigenetic signal given that IGF1 is a key determinant of growth and aging (Junnila et al., 2013; Laron, 2001).

### Construction of an epigenetic clock in sheep

We then established epigenetic clocks from our sheep blood and ear methylation data, respectively, as well as a combined blood and ear clock (hereafter referred to as the *multi-tissue clock*) using a penalized regression model (elastic-net regression). In total, 185 CpG sites were included for the multi-tissue clock, which was shown to have a MAE of 5.1 months and an age correlation of 0.95 when calculated using a leave-one-out cross validation (LOOCV) analysis (Figure 2A). Taking into account the expected lifespan of sheep in commercial flocks (10-12 years), the error of the multi-tissue clock is 3.5-4.2 % of the lifespan – comparable to the human skin and blood clock at ∼3.5 % (Horvath et al., 2018) and the mouse multi-tissue clock at ∼5 % of expected lifespan (Meer et al., 2018; Petkovich et al., 2017; Stubbs et al., 2017; Thompson et al., 2018; Wang et al., 2017).

**Figure 2.**
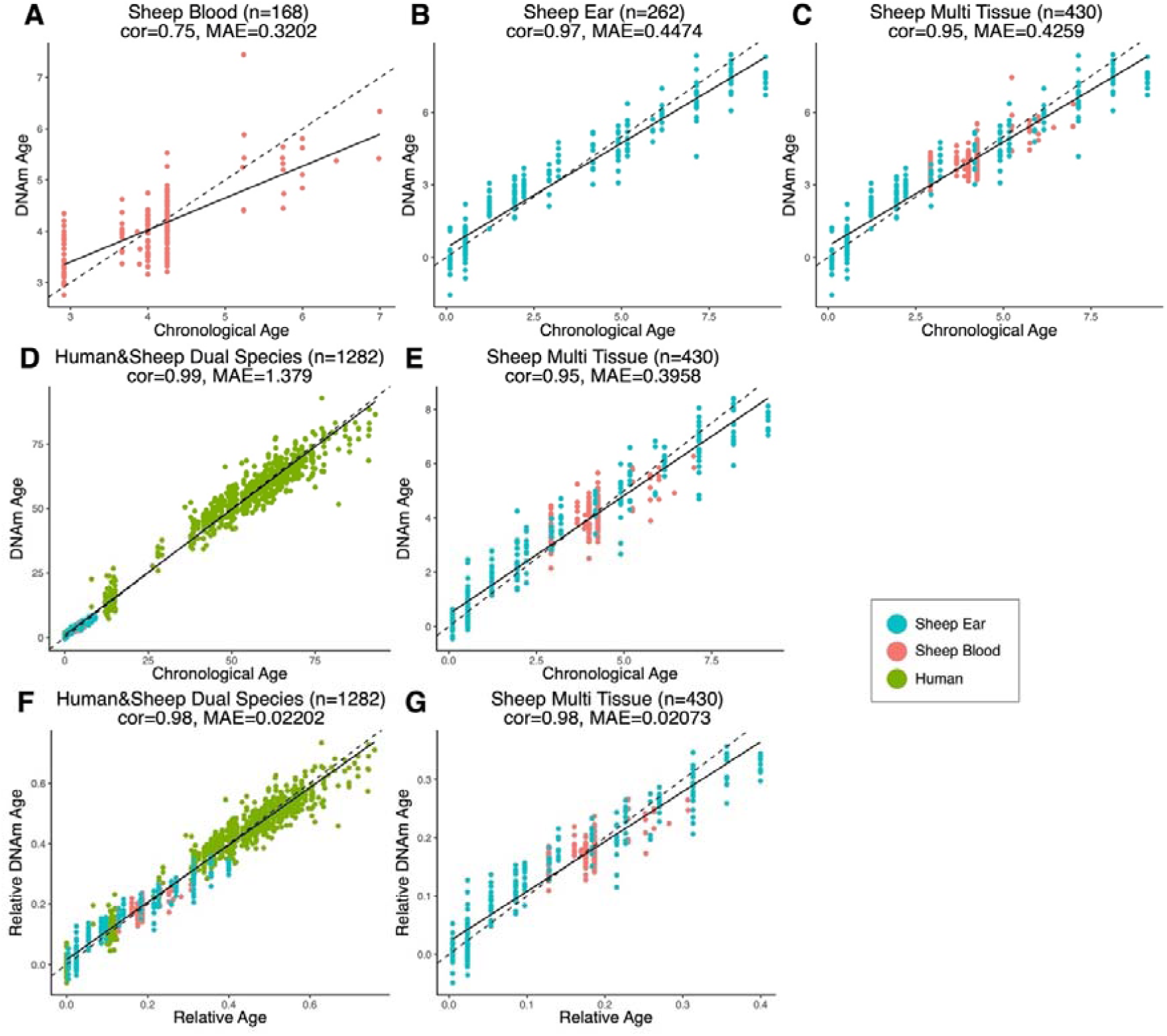
Comparison of chronological age (x-axis) and epigenetic age (y-axis) for a variety of clock models. Correlation (cor) and median absolute error (error) is indicated for **A)** sheep blood (cor=0.75, error=0.3202), **B)** sheep ear (cor=0.97, error=0.4474), **C)** sheep multi-tissue (ear and blood) (cor=0.95, error=0.4259), **D)** human & sheep dual-species (cor=0.99, error=1.379), and **E)** sheep multi-tissue (cor=0.95, error=0.3958). **F)** chronological age (x-axis) plotted against epigenetic age (y-axis) relative to maximum lifespan for human & sheep dual-species clock (cor=0.98, error=0.02202), and **G)** chronological age (x-axis) plotted against epigenetic age (y-axis) relative to maximum lifespan for the sheep multi-tissue clock (cor=0.98, error=0.02073). Maximum lifespan values used were for human and sheep respectively were 122.5 years and 22.8 years. Each data point represents one sample, coloured based on origin.

Two human and sheep dual-species clocks were also constructed, which mutually differ by way of age measurement. One estimates the chronological ages of sheep and human (in units of years), while the other estimates the relative age – a ratio of chronological age of an animal to the maximum known lifespan of its species (defined as 22.8 years and 122.5 years for sheep and human respectively) with resulting values between 0 and 1. The measure of relative age is advantageous as it aligns the ages of human and sheep to the same scale, yielding biologically meaningful comparison between the two species. The dual-species clock for chronological age leads to a median error of 16.53 months when considering both species, or 4.74 months for sheep only (Figure 2D-E). The dual-species clock for relative age produced median errors of 0.020 of the maximum lifespans for both species (approximately 2.45 years for human, or 5.4 months for sheep) and 0.021 for sheep only (approximately 5.7 months) (Figure 2F-G).

### Castration delays epigenetic aging in sheep

To test the role of androgens in epigenetic age acceleration, we exploited the fact that castrated male Merino sheep are frequently left to age ‘naturally’ on New Zealand high-country farms in return for yearly wool production, in contrast to non-breeding males of other sheep varieties which are usually sold as yearlings for meat. Both castrated males and intact aged-matched controls were sourced from genetically similar flocks kept under comparable environmental conditions. Interestingly, both intact and castrated males showed equivalent epigenetic age during the juvenile years, however, once they advanced beyond the yearling stage, castrates appeared to have slowed rates of epigenetic aging (Figure 3A). Indeed, when we only considered sheep beyond 18 months of age, we found castrates had significantly reduced epigenetic age compared to intact male controls (Figure 3B, p=0.018). While the extent of the age deceleration consistently increased with advancing age, mature castrates were on average epigenetically 3.1 months ‘younger’ than their chronological age (Figure 3B). In contrast, DNAm age of intact males was comparable to their chronological age (0.14 months age decelerated), as were females (0.76 months age accelerated), who comprised the majority of the samples from which the clock was constructed. Notably, the age deceleration observed in castrates was corroborated using the human & sheep dual-species clock (Figure S3, p=0.04).

**Figure 3.**
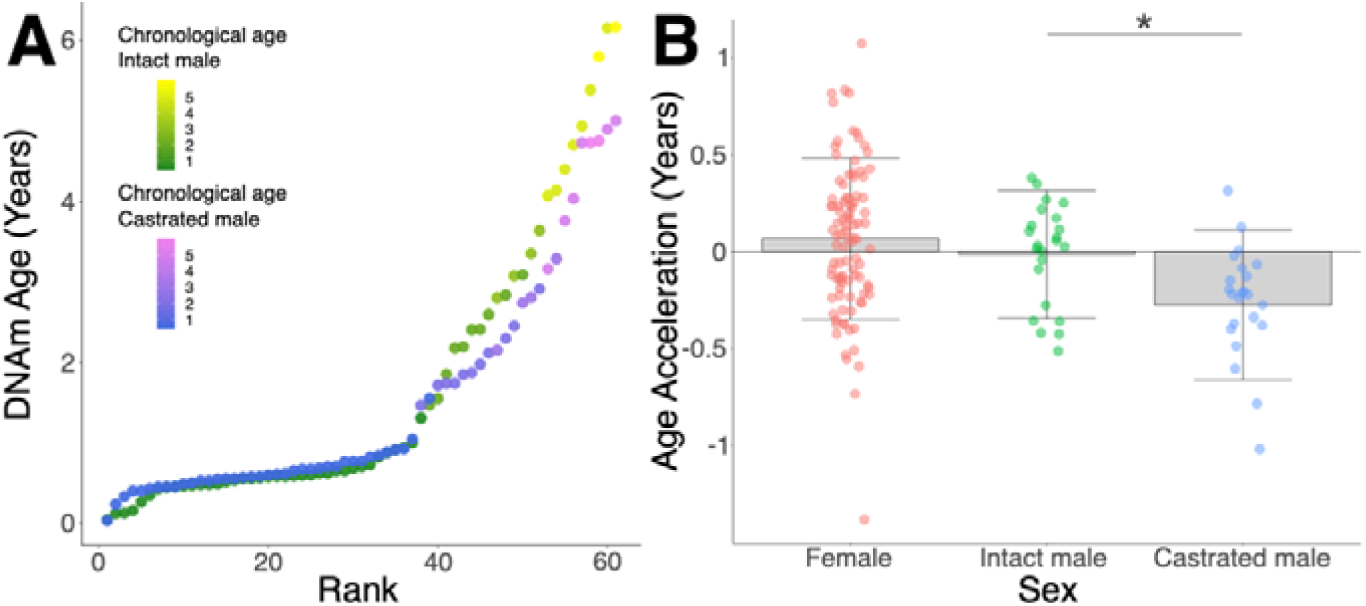
Epigenetic age deceleration in castrated sheep. **A**) Epigenetic age in age-matched castrated and intact males. To equate the cohort sizes for intact and castrated males, two age-matched castrates with DNAm age estimates closest to the group mean were excluded. **B**) Age acceleration based on sex and castration status in sexually mature sheep only (ages 18 months+ only). Castrated males have decelerated DNAm age compared to intact males (*: p=0.01, Mann-Whitney U test).

To explore the mechanistic link between androgens and epigenetic aging, we identified probes with significant differences between the rate of age-dependent methylation changes in castrated or intact males (Table S3). We found there was a sharp inflection in *p* value after approximately the 50 most significant probes (Figure S4A) and thus represented a natural cut-off for analysis. A recent comparison of age-related methylation changes in the blood of human males and females revealed that almost all regions of interest appeared to be X-linked (McCartney et al., 2019). Given that there are already well characterised differences between male and female methylation patterns on the X-chromosome as a result of gene-dosage correction (Heard and Disteche, 2006), it could be argued that X-linked age-related differences may be driven by peculiarities of methylation arising from X-chromosome inactivation, as opposed to differences in androgen production *per se*. To test this for sheep, we examined the genomic location of our asDMPs, and found they are evenly distributed between individual autosomes and the sex chromosomes (Figure S4B-C).

Interestingly, we found several sites that become progressively hypomethylated in intact males with age but maintain a consistent level of methylation throughout life in castrates and females (Figure 4A-D). Indeed, of the top 50 most significantly different asDMPs, only two (cg03275335 *GAS1* and cg13296708 *TSHZ3*) exhibited alteration whereby intact males gained methylation (Figure 5A). We found that many asDMPs we identified were linked to genes known to be regulated by androgen receptor (AR) (e.g. *MKLN1, LMO4, FN1, TIPARP, ZBTB16* (Jin et al., 2013)) and as such, we were encouraged to find further mechanistic connections between asDMPs and TF regulation.

**Figure 4.**
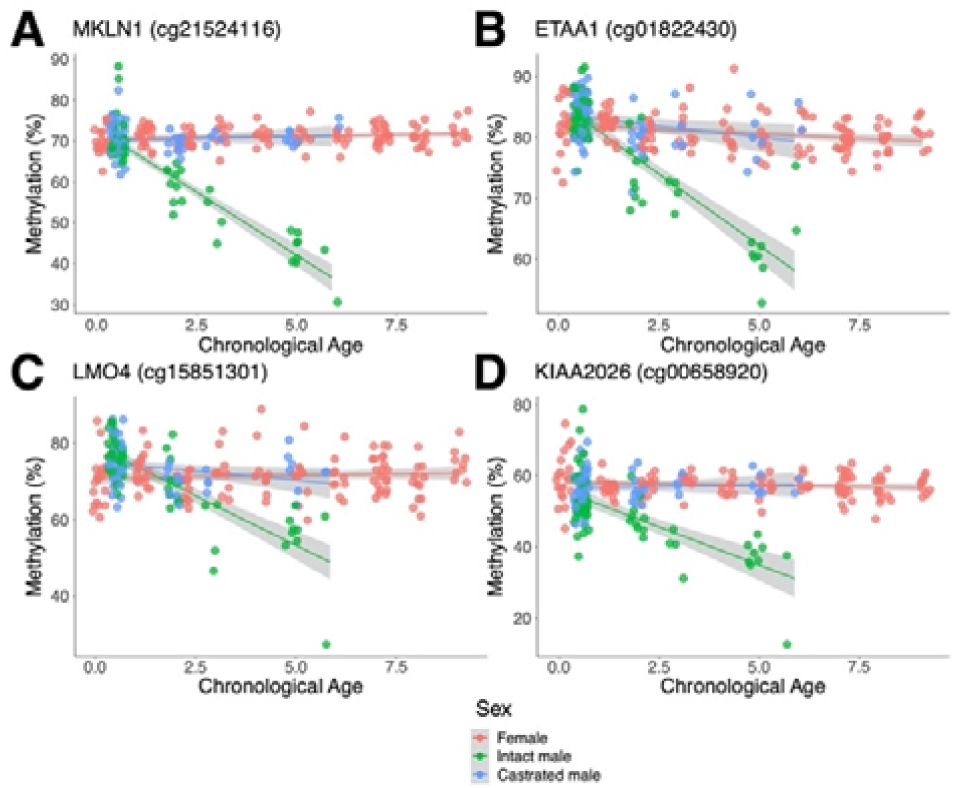
Top androgen-sensitive differentially methylated probes (asDMPs) in sheep ear. **A)** *MKLN1* (cg21524116, p=1.05E^-27^), **B)** *ETAA1* (cg01822430, p=1.31E^-13^), **C)** *LMO4* (cg15851301, p=1.62E^-09^), **D)** *KIAA2026* (cg00658920, p=2.46E^-09^). The *p* values were calculated using a t-test of the difference in linear regression slopes.

**Figure 5.**
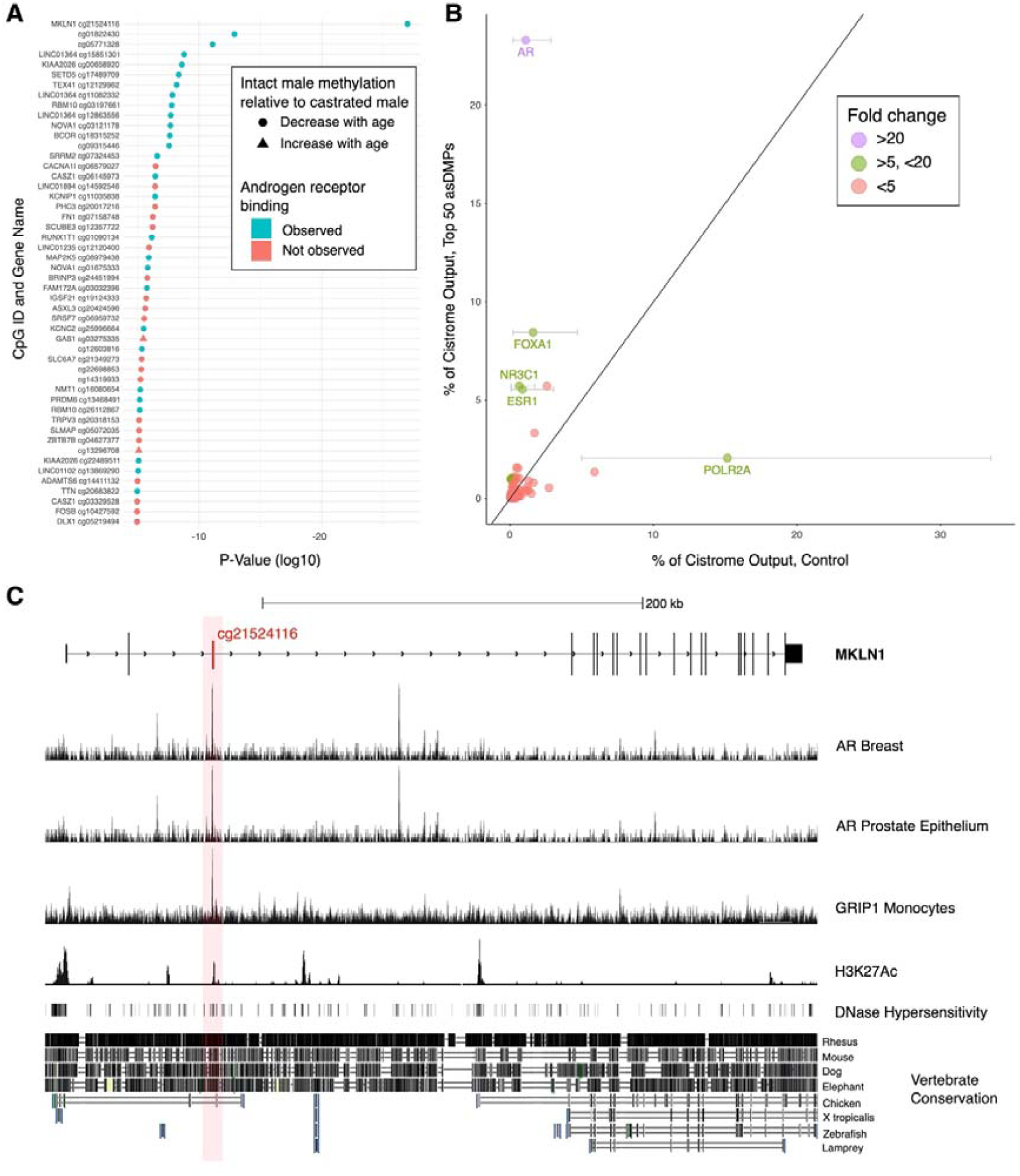
Analysis of chromatin immunoprecipitation and sequencing (ChIP-seq) data indicates functional links to sex-dependent epigenetic aging. **A)** Top 50 asDMPs between intact and castrated male sheep, and the human genes they map to (where applicable). The top 14 most significant asDMPs are all bound by AR; and 48/50 of these sites exhibit hypomethylation with age in intact males relative to castrated males. **B)** Observed transcription factor binding at the top 50 asDMPs compared to expected binding based upon empirical sampling at random CpGs (average of 1000 bootstrap replicates). Transcription factors with >5-fold variation and an absolute value of >2% are labelled with error bars showing the range of TF binding in bootstrap sampling. Colours indicate fold-change between observed and expected TF binding; 1-5 (red), 5-20 (green) and >20 (purple) **C)** Genomic view of *MKLN1* containing the most significant asDMP cg21524116 illustrating AR binding and indicators of active chromatin.

To do this, we used examined TF-binding of the human regions homologous to our asDMPs using the Cistrome Data Browser Toolkit - although Cistrome contains data from a wide-range of transcription factors, we noted AR binds to over half the asDMPs (28/50), with the top 14 most significant asDMPs all showing AR binding (Figure 5A). To ensure that this was not a result expected by chance alone, we performed empirical sampling whereby 1000 replicates of the identical analysis was performed but with 50 random CpG sites from the methylation array at each bootstrap replicate. The observed/expected enrichment for AR binding in our 50 asDMPs was 22-fold (23.29%/1.09%, p=<0.001), and much higher than enrichment for any other TF. Nevertheless, there were several other interesting related TFs with high observed/expected enrichment for asDMPs, including the estrogen receptor (ESR1), the glucocorticoid receptor (NR3C1) and forkhead box A1 (FOXA1) (Figure 5B).

While we saw similar features at other asDMPs (S5A-B), the asDMP that was the most different in epigenetic aging rate between castrates and males, *MKLN1*, stood out as being particularly interesting from a gene regulatory perspective (Figure 5C). Overlapping with this site, and AR binding, were peaks of DNase I hypersensitivity, H3K27ac histone marks as well as good vertebrate conservation compared to surrounding sequences.

### Androgen-sensitive DMPs are present in divergent mammalian species but are tissue specific

To determine if the androgen-dependent methylation exists in divergent mammalian groups, we assessed methylation changes at these asDMPs in mouse tissues (blood, cerebellum, cortex, liver, muscle and striatum). Again, cg21524116 in *MKLN1* stood out - mouse muscle exhibited the same sex-specific trend in as seen in sheep, whereby females retain a constant level of high methylation was retained in females while methylation levels gradually decreased with age in males (p=0.003) (Figure 6A). However, this same trend could not be seen in mouse (Figure 6B) or sheep blood, suggesting that this androgen-sensitivity is tissue-specific.

**Figure 6.**
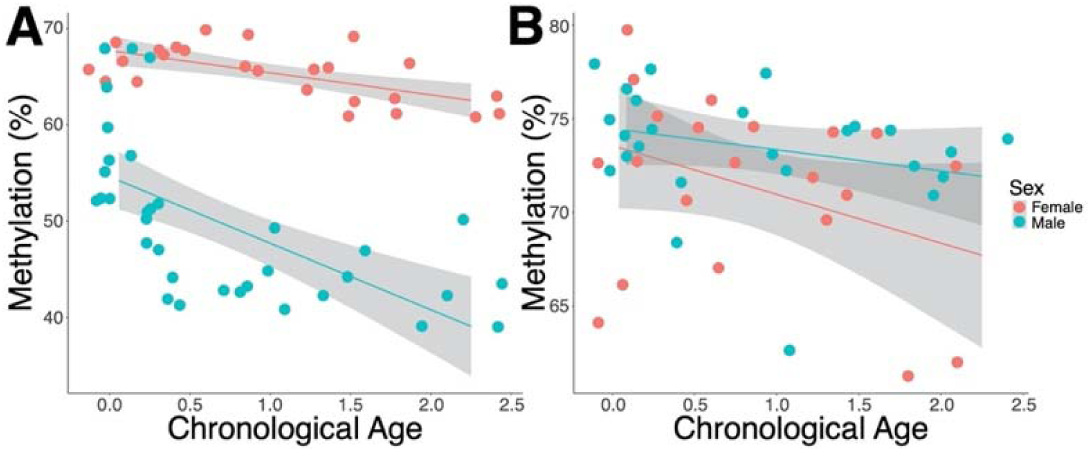
Androgen-sensitive methylation patterns in cg21524116 (*MKLN1*) in mouse muscle and blood. **A)** *MKLN1* in muscle of male mice exhibits a similar androgen-dependent methylation loss as seen in intact sheep, suggesting the existence of a wider mammalian effect (p=0.0035, t-test). **B)** In contrast to muscle, *MKLN1* in mouse blood does not demonstrate androgen-sensitivity (p=0.362, t-test), indicating that this effect is tissue-specific. The other tissues examined (cerebellum, cortex, liver and striatum) also showed no androgen-sensitive methylation patterns.

## DISCUSSION

Epigenetic clocks are accurate molecular biomarkers for aging which have proven to be useful for identifying novel age-related mechanisms, diseases, and interventions that alter the intrinsic aging rate (Horvath and Raj, 2018). Here, we developed the first epigenetic clock for sheep and show it is capable of estimating chronological age in sheep with a MAE of 5.1 months – between 3.5 % and 4.2 % of the average sheep lifespan. Significantly, we also present the first evidence that castration feminises parts of the epigenome and delays epigenetic aging.

Improved survival has previously been reported in castrated sheep compared to intact males and females, part of which was suggested to be attributed to behavioural changes such as reduced aggression (Jewell, 1997). Our data shows that castration also causes a delay in intrinsic aging as assessed by the epigenetic clock, with an average reduction in epigenetic age of 3.1 months (Figure 3B). Moreover, delayed epigenetic aging in castrates is also seen relative to intact males and females, which is consistent with castrated males outliving intact animals of both sexes (Jewell, 1997). We also find that the degree of age deceleration observed in castrated males is dependent on their chronological age. For instance, the average DNAm deceleration is increased by an additional 1.2 months when considering individuals aged beyond 2.9 years. In contrast, we saw no difference between castrated and intact males younger than 18 months. Together this implies that the effects of androgen exposure on the epigenome and aging are cumulative. Similar findings of greater age-deceleration at later chronological ages have been observed in rodent models, with long-lived calorie-restricted mice showing a younger epigenetic age relatively late in life, but similar epigenetic aging rates at younger ages (Petkovich et al., 2017).

These results support the reproductive cell-cycle theory as an explanation for sex-dependent differences in longevity of mammals (Atwood and Bowen, 2011; Bowen and Atwood, 2004). Androgens and other testicular factors may be working in an antagonistic pleiotropic manner whereby they push cells through the cell cycle and promote growth in early life to reach reproductive maturity, thus also influencing the epigenome in an age-related manner. This process, however, may become dysregulated and promote senescence at older ages, reflected in the hastening of the epigenetic clock observed in intact males compared to castrates. It is well-known in farming practise that where it can be managed appropriately, leaving male sheep intact or partially intact (i.e. cryptorchid) increases body mass (Seideman et al., 1982),something we also observed in our study (Figure S6). This indicates greater rates of cell cycle progression, cellular division, and tissue hyperplasia. Under this hypothesis, the effects of castration should depend on whether animals are castrated before or after puberty. In rats, castration just after birth (i.e. prior to puberty) causes substantial lifespan extension while castration after puberty has smaller effects (Talbert and Hamilton, 1965), supporting the idea that male gonadal hormones have effects at early life stages that have deleterious consequences for survival.

Consequences of castration for increased survival and slowed epigenetic aging could also be linked to the effects of androgens on sexual dimorphism and adult reproductive investment (Brooks and Garratt, 2017). Life-history theories predict that males in highly polygynous species, like sheep, should be selected to invest heavily in reproduction early in life, even at the expense of a shorter lifespan, because they have the potential to monopolise groups of females and quickly produce many offspring (Clutton-Brock and Isvaran, 2007; Tidiere et al., 2015). By contrast, selection on females should promote a slower reproductive life strategy, because female reproductive rate is limited by the number of offspring they can produce. While we show that castration slows epigenetic aging in sheep, loss of ovarian hormone production in mice and human (through ovariectomy or menopause) is associated with a hastening of the epigenetic clock (Levine et al., 2016; Stubbs et al., 2017), consistent with the beneficial effects of female ovarian hormones on survival. Thus, it appears that both male and female sex-hormones differentially regulate the epigenetic aging process in directly opposing ways, in a manner that is consistent with the life-history strategies classically thought to be optimal for each sex.

Our results provide further insight into the mechanisms of aging and genes affected by age-associated methylation. Several well-described growth and transcription factors demonstrate a high positive age correlation; including *FGF8*, a developmental growth factor involved in embryonic brain and limb formation (Lorenzi et al., 1995; MacArthur et al., 1995). Low methylation at this site in our youngest samples may be indicative of some residual expression following birth, which is quickly silenced by post-natal hypermethylation (Figure 1D). In females, genic *IGF1* methylation demonstrates the opposite trend. Methylation levels observed at position cg18266944 in intron 2 of *IGF1* are highest immediately after birth followed by rapid demethylation, consistent with an activation of the gene to promote post-natal growth (Baker et al., 1993). Given that *IGF1* and its associated mitogenic pathway is one of the most widely studied molecular driver of aging, we considered this a relevant discovery warranting further exploration. Despite this site being intronic, considerable sequence conservation and its position within a DNase hypersensitivity site indicate that this locus may have some regulatory function (Figure S5C), however, it is not possible to determine if the same process occurs in males due to a lack of equivalent early life samples. Importantly, peaks of AR binding have been observed immediately upstream of the *IGF1* gene, which could suggest some transcriptional control of *IGF1* by androgens. Rising methylation levels in DNA detecting by probes mapped to other well-characterised transcription factors, *PAX5, PAX6*, and *HOXC4*, are indicative of larger transcriptional shifts over the lifespan (Figure 1C), concordant with the results from the gene ontology analysis (Table S2).

Comparison of intact and castrated males also allowed us to identify several age-related DMPs that display clear androgen-sensitivity (Figure 4A-D), with castrated males exhibiting a feminised methylation profile compared to intact counterparts. In contrast to similar experiments performed in humans, we found that these sex-specific CpG sites are not predominantly X-linked in sheep (McCartney et al., 2019), but are instead distributed evenly throughout the genome (Figure S4). The most striking example of age-dependent androgen-sensitive methylation loss is that detected by the probe cg21524116, mapping to *MKLN1* (Muskelin) (Figure 4A). Sex-specific methylation is also seen at this probe location in mouse muscle but in neither sheep or mouse blood (Figure 6A-B), implying that such androgen-sensitive effects in *MKLN1* and other loci may be mammalian wide but certainly not ubiquitous across all tissues. Evidence for *MKLN1* androgen-dependency has previously been presented (Jin et al., 2013) and MKLN1-containing complexes have been shown to regulate lifespan in *Caenorhabditis elegans* (Hamilton et al., 2005; Liu et al., 2019), although no links between this gene and mammalian longevity have yet been reported. ChIP-seq data demonstrates enriched AR binding at the position of this asDMP in human, as well as exhibiting high sequence conservation, DNase hypersensitivity and H3K27ac marks – the latter two of which are markers of open chromatin and indicate transcriptionally active areas (Figure 5C) (Creyghton et al., 2010; Wang et al., 2008). Taken together, this evidence implies that the site we identified in *MKLN1* is a reliable biomarker of androgen-induced aging in sheep, and it may also have some regulatory function involved in male-accelerated aging.

Similarly, many of the other highly significant asDMPs (58%) are also bound by androgen receptor (Figure 5A). When considering the TF binding compared to background levels, our data shows that particularly AR, but also ESR1, FOXA1 and NR3C1 binding are enormously enriched in our top asDMPs; all of which share biologically integrated functions. NR3C1, which encodes the glucocorticoid receptor (GR) and has been previously linked to longevity in certain populations (Olczak et al., 2019), is an anabolic steroid receptor and thus shares significant homology in its binding domain and targets many DNA sequences also bound by AR (Claessens et al., 2017). FOXA1 has been found to regulate estrogen receptor binding (Carroll et al., 2005; Hurtado et al., 2011) as well as AR and GR binding (Sahu et al., 2013) in both normal and cancer cells. Furthermore, FOXA1 aids in ESR1-mediated recruitment of GRs to estrogen receptor binding regions (Karmakar et al., 2013). Interestingly, AR agonist treatment in breast cancer models reprograms binding of both FOXA1 and ESR (Ponnusamy et al., 2019) suggesting some degree of antagonistic function between the androgen and estrogen receptors. If this is true for asDMPs in sheep, these sites may well represent a conduit through which castrates take on physiologically feminised traits, including delayed aging.

Having said this, it remains a possibility that methylation levels at these androgen-sensitive sites has very little to do with biological aging and instead only diverge as time progresses due to the period of androgen exposure or deficiency. Specifically, the changes in methylation observed between intact and castrated males may not be adaptive at all, and rather, methylation is progressively “diluted” by binding of androgen receptor to the DNA. Variable methylation at AR target genes has been reported in humans with androgen insensitivity syndrome (AIS) when compared to normal controls, supporting the notion that AR binding influences methylation at target genes (Ammerpohl et al., 2013). However, the authors noted that these methylation shifts were sporadic, which does not explain the consistent feminisation of methylation levels we observe in castrated sheep at many asDMPs.

As yet, we do not know if castration in later life would drive feminisation of methylation patterns as we observed for early life castration (Figure 4). This is however, an interesting consideration - it is possible that castration late in life would quickly recapitulate the methylation differences seen in those castrated early in life, or it may be that methylation patterns established during early growth and development are difficult to change once set on a particular aging trajectory. This distinction may be important from a functional perspective because early and later-life castration can have differing effects on survival in rodents (Talbert and Hamilton, 1965). Moreover, while early-life castration has been shown to extend human lifespan, androgen depletion in elderly men can be associated with poor health (Araujo et al., 2011).

In summary, this paper describes a robust epigenetic clock for sheep that is capable of estimating chronological age, detecting accelerated rates of aging, and contributes to a growing body of work on epigenetic aging. In addition to demonstrating the utility of sheep as an excellent model for aging studies, our data identify androgen-dependent age associated methylation changes that affect known targets of sex hormone pathways and hormone binding transcription factors. While these changes may not promote aging *per se*, identification of loci with age-dependent androgen-sensitive methylation patterns uncovers novel mechanisms by which male-accelerated aging in mammals can be explained.

## Acknowledgements

We thank Ari Samaranayaka for his guidance with portions of the statistical analyses.

## SUPPLEMENTARY FIGURES

**Supplementary Figure 1.**
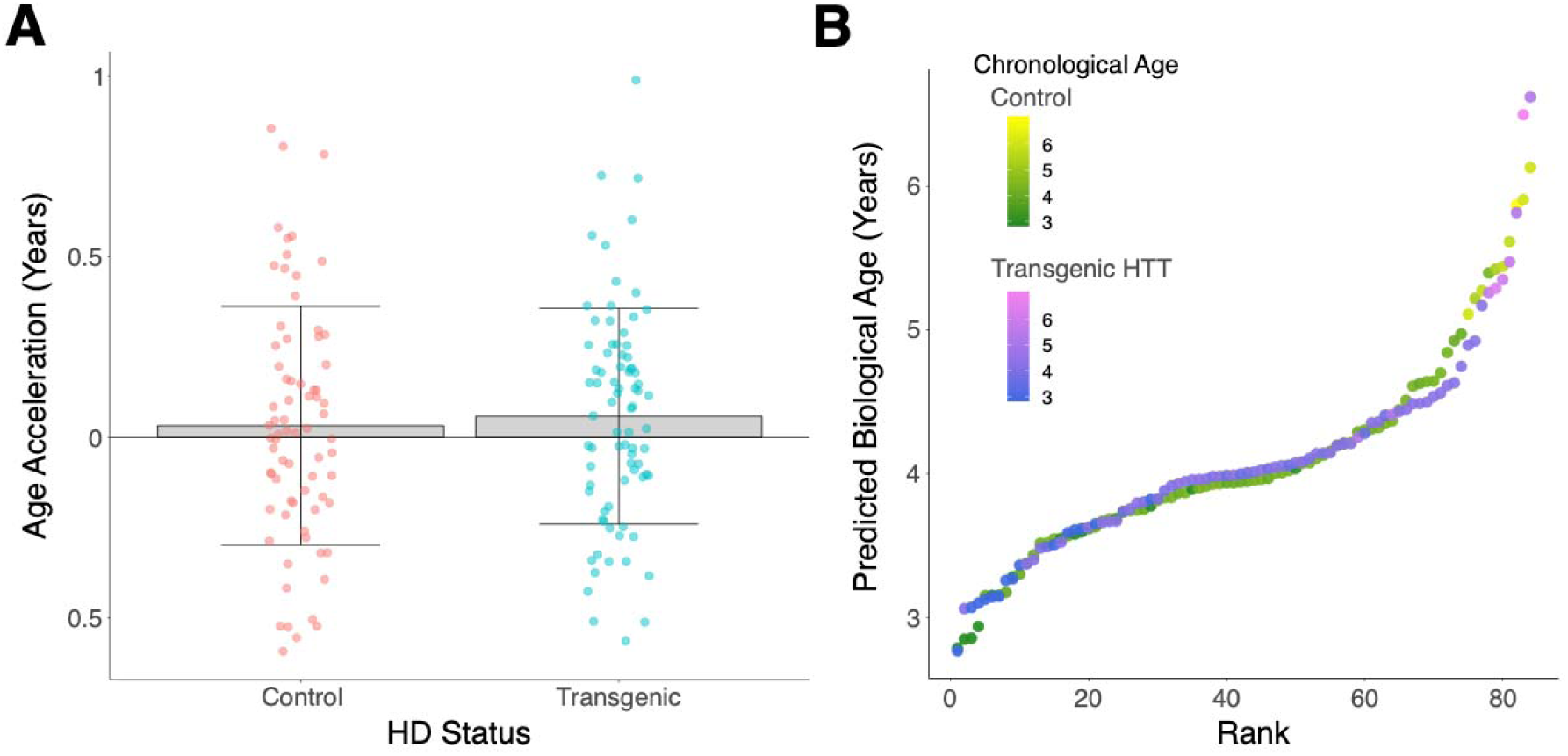
The epigenetic ages of control and transgenic HTT sheep are not significantly different (p=0.30, Mann-Whitney U test).

**Supplementary Figure 2.**
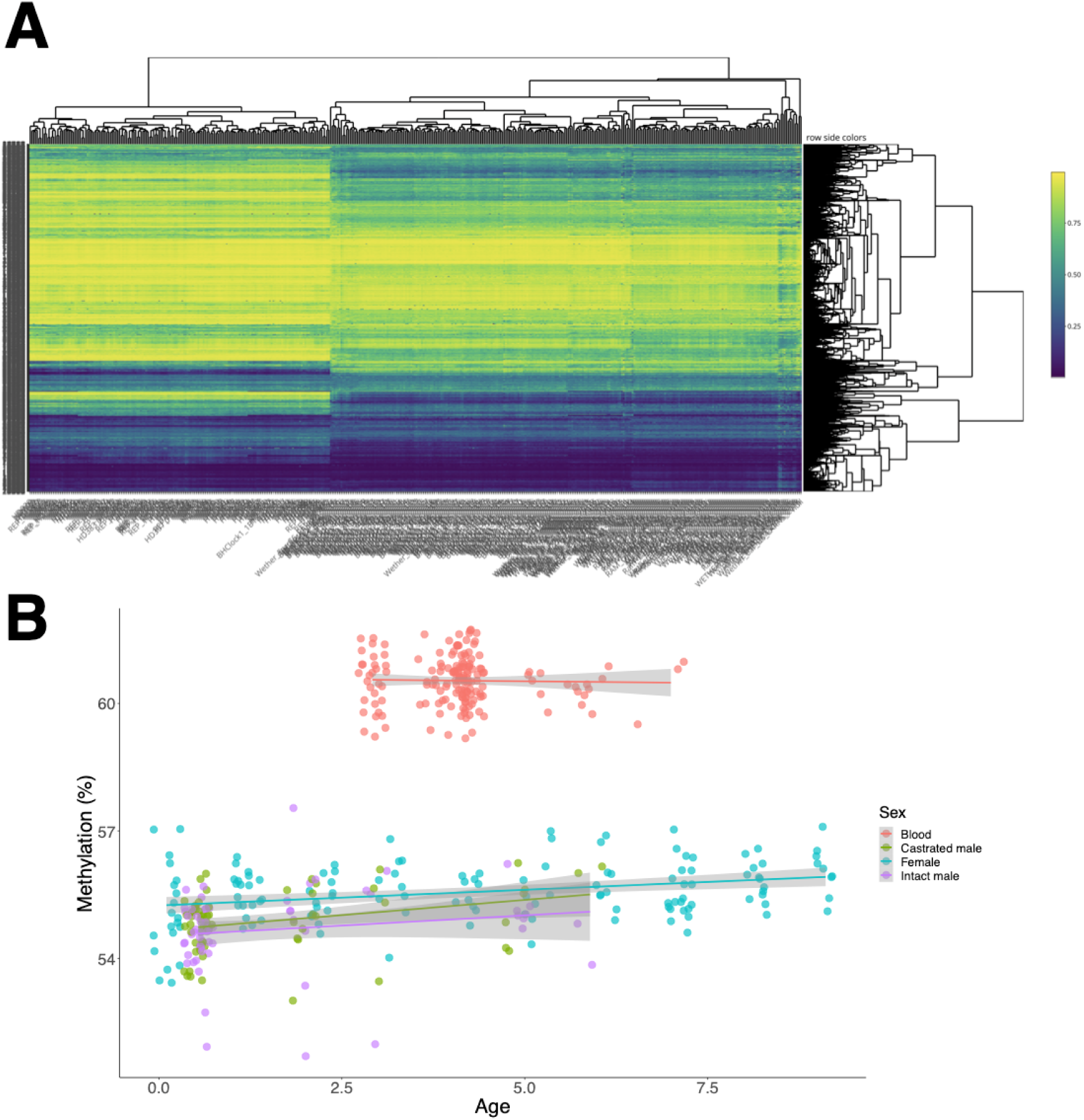
**A)** Hierarchical clustering heat map shows methylation profiles grouped by sheep tissue type; blood (left side) vs. ear (right side). **B)** Average methylation in sheep ear and blood as age progresses.

**Supplementary Figure 3.**
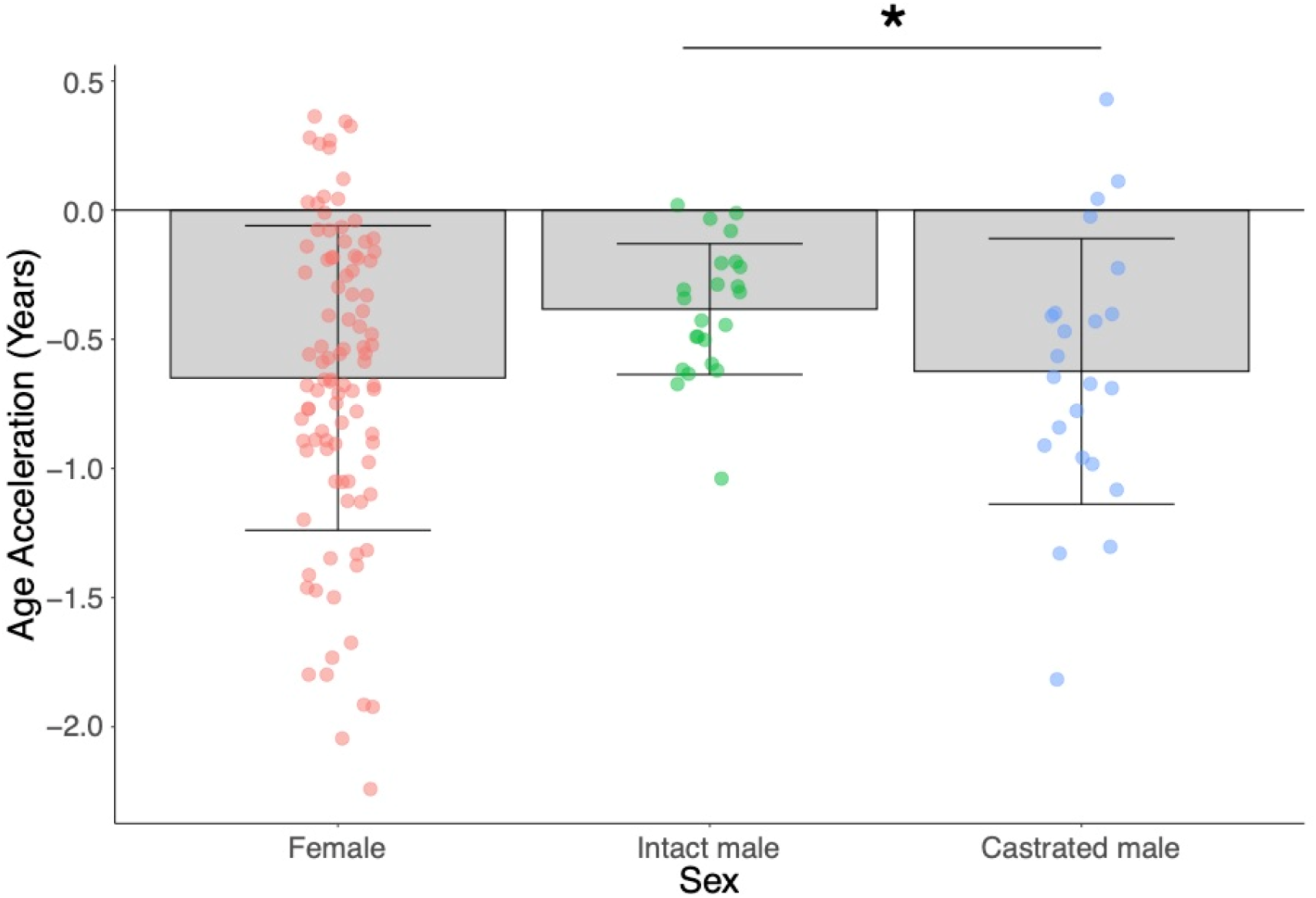
Age acceleration based on sex and castration status in sexually mature sheep only (age 18 months+ only), using the human & sheep dual-species clock. Asterisk indicates the significant difference between age acceleration in castrated and intact males (p=0.04, Mann-Whitney U test).

**Supplementary Figure 4.**
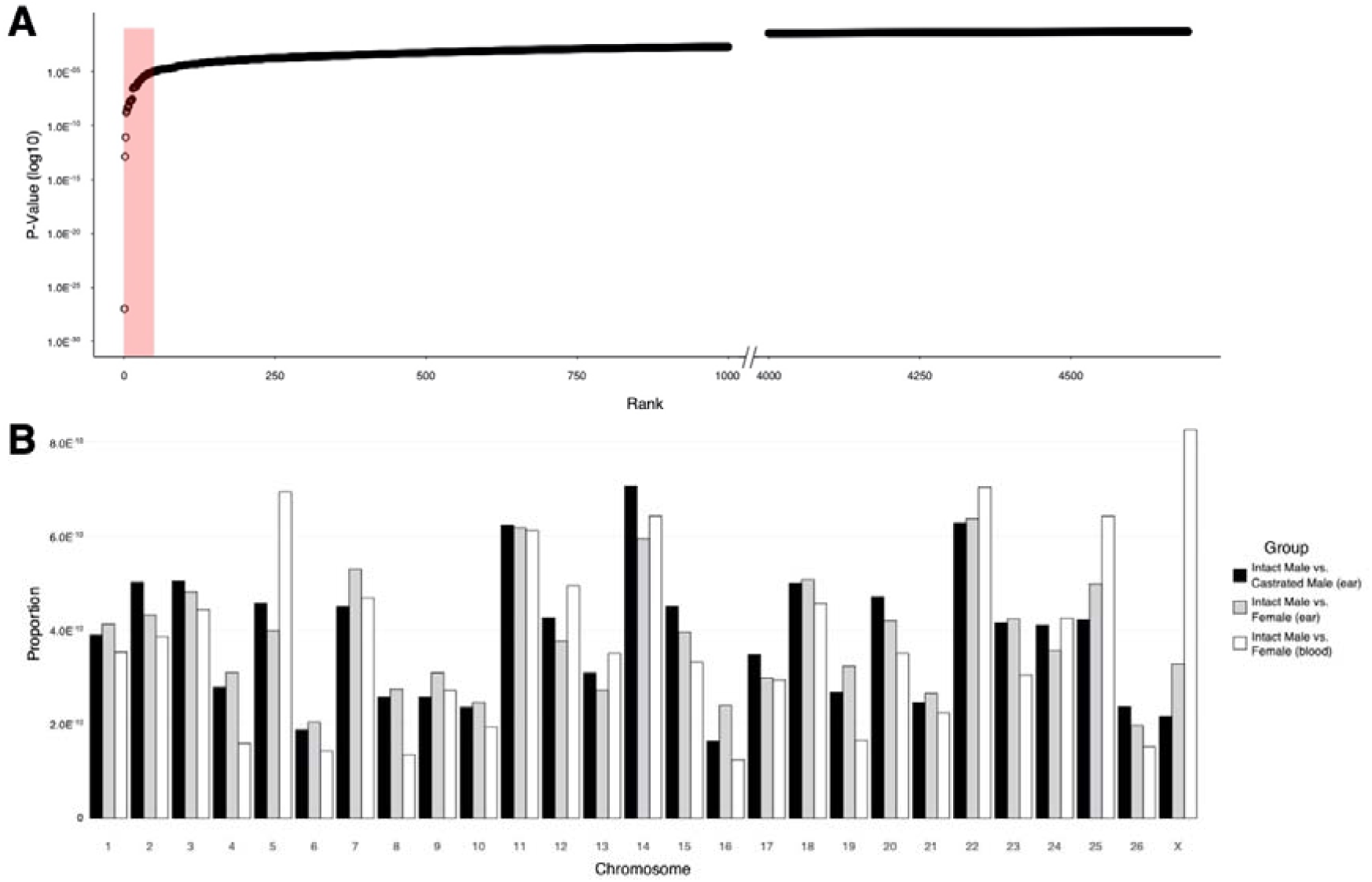
Chromosome location of androgen-sensitive DMPs. **A)** All 4,694 statistically significant (p<0.05) asDMPs ordered by *p* value. The top 50 asDMPs are highlighted in red (and are examined more closely in Figure 5A), after which there is a clear inflection point of *p* value. **B)** Chromosome location of all statistically significant asDMPs in all sheep groups. Y-axis shows the proportion of probes that map to each chromosome, normalised for chromosome size and percentage of significant probes within each comparison group.

**Supplementary Figure 5.**
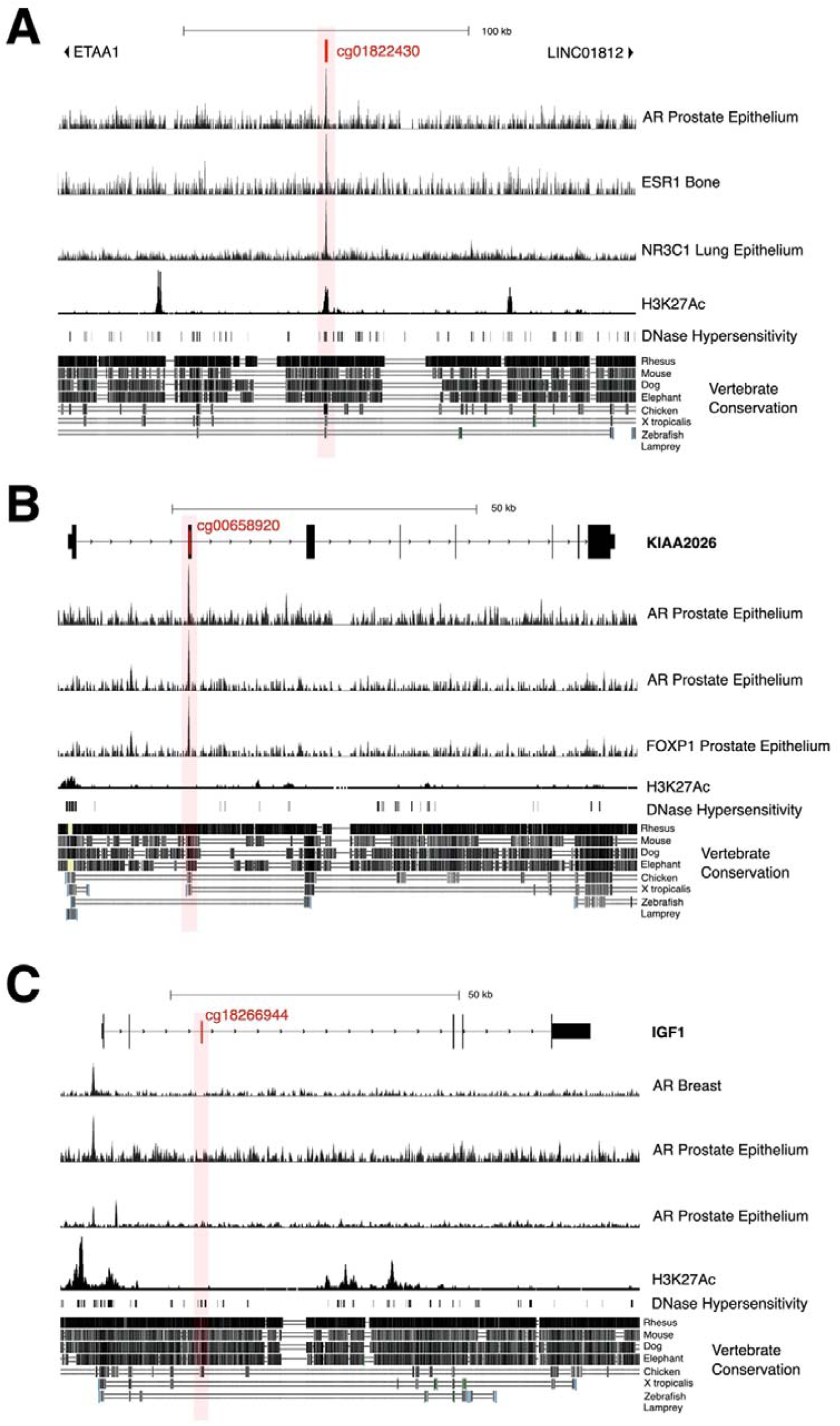
Gene views of key androgen-sensitive sites showing evidence for possible regulatory functions. **A)** *ETAA1*; cg01822430 (second most significant sheep asDMP). **B)** *KIAA2026* cg00658920. **C)** *IGF1* cg18266944.

**Supplementary Figure 6.**
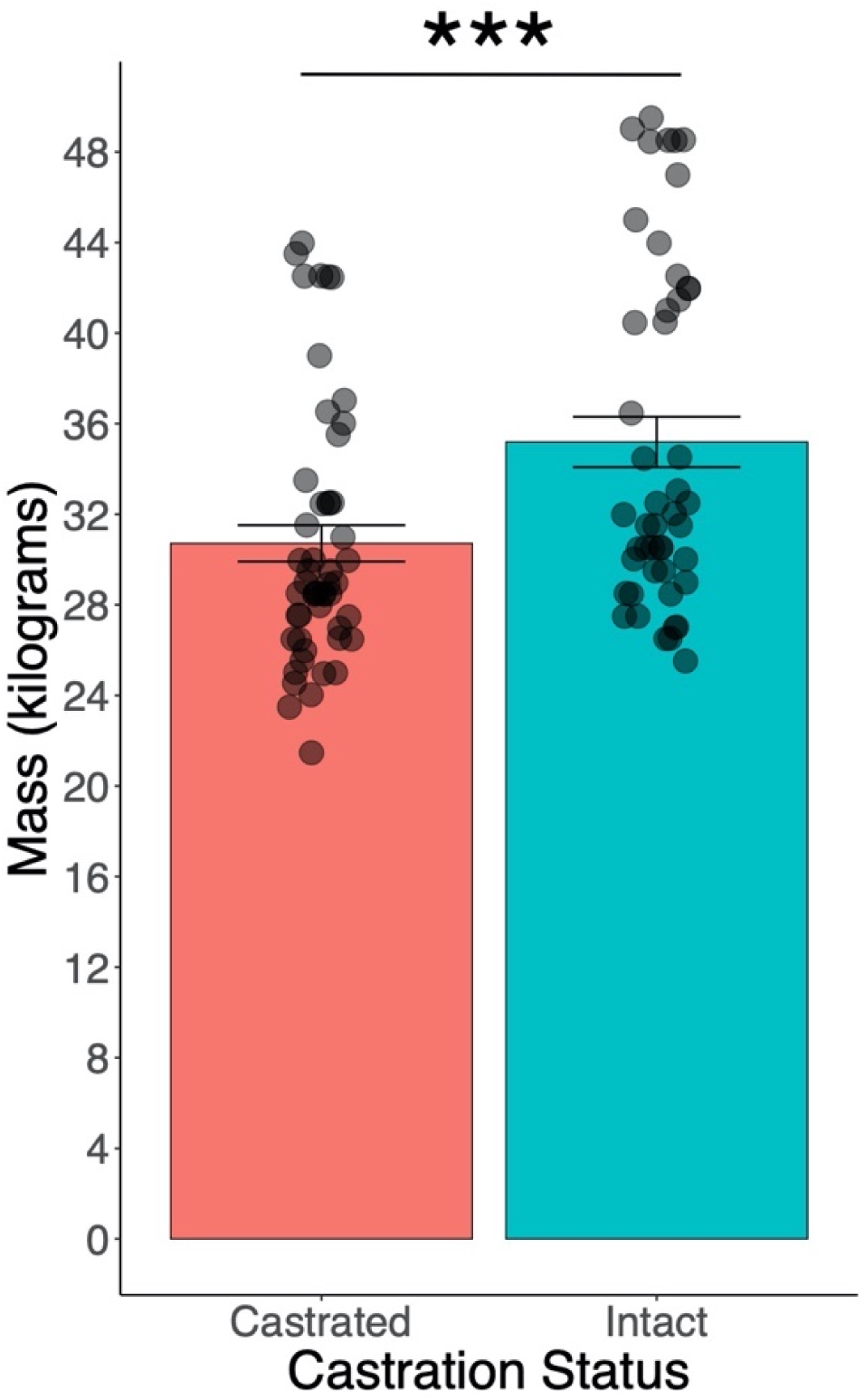
Mass (kg) of male lambs (<1 year) dependent on castration status. p=<0.001 (T-test). Error bars = SEM.

## Notes

### Competing Interest Statement

SH is a founder of the non-profit Epigenetic Clock Development Foundation which plans to license several patents from his employer UC Regents. These patents list SH as inventor. TAH and DMB are a shareholders and directors of Totovision Ltd, a small agricultural and biotechnology consultancy. The other authors declare no conflicts of interest.

